# Novel methods of isolation and amplification of progenitor cells applied to avian primordial germ cells

**DOI:** 10.1101/108571

**Authors:** Mariacruz Lopez-Diaz

**Author notes:** Present address: Instituto Nacional de Investigación y Tecnología Agraria y Alimentaria (INIA). Departamento de Reproducción Animal. 28040 Madrid – Spain.

## Abstract

**Background:** The progenitor cells in adult tissues are scarce and have a great regenerative potential. In this study novel methods were used to improve the isolation and culture of the chicken primordial germ cells (PGCs) from stage X and HH 8-9 embryos. The cellular size and external glycoprotein envelope were the two criteria studied and used.

**Results:** PGCs were segregated with high efficiency and purity, from stage X and HH 8-9 gross cell suspensions through cell strainers with 10 μm of pore size. In embryos *in toto, WGA* Alexa 594 (affinity for N-acetylglucosamine) and *Con A* Alexa 488 (binding D-mannosyl) were used to characterize external polysaccharides of the PGCs. The PGCs in stage X embryos (zone pellucida), have predominately Nacetylglucosamine and later on, in HH 8-9 embryos (cephalic zone), α-D mannosyl residues, in a specific manner. In coated plates with the appropriate lectin and in alkaline conditions, isolated cells from stage X and HH 8-9 embryos formed numerous clumped PGC-LCs with spherical shape “germspheres”. In all isolates from single embryo, immunohistochemistry confirmed that they were PGCs and revealed that the “germspheres” were formed by hundreds of positive cells to *VASA* and *SSEA-1*. N-acethyl D+glucosamine supplementation to the culture media greatly enhances the amplification of isolated PGC-LCs.

**Conclusions:** These gentle and quick strategies with high yields of PGCs can be potentially useful for many progenitor cells in Regenerative Medicine.

## BACKGROUND

Just like any other progenitor cells in adult tissues, primordial germ cells (PGCs) are scarce during embryonic development. This is the main reason why in mammals the culture of PGCs, precursors of oocyte and spermatozoa, can only be achieved for short periods. Althought sophisticated strategies are being developed to circumvent this inconvenient, like 3D engineering biotechnologies to imitate the germ cell niche (1) or the complicated differentiation procedures of embryonic stem cells (ESCs) or induced pluripotent stem cells (iPSC) into PGCs, still PGCs are not optimally sustained (2).

Things are quite different for avian PGCs whose origin, morphology and migration have been well characterized (3, 4, 5) since first described by Swift (6). Several advantages have been pointed out in the avian embryo: the development can be openly observed and the PGCs migrate to the gonads throught the blood stream from where they can be isolated. When thousands of PGCs are in blood or have colonized the gonads (gonocytes) many procedures have been described for the enrichment of the cell suspensions, from unspecific ficoll separation (7, 8) to efficient IMACS or FACS sorting (9), yet yields of germ cells are commonly low and their manipulation is tricky. Besides, the methodologies suffer a consistent gender bias: most culture systems have succeeded with male PGCs extracted from blood (10, 11,12), but only in few instances could female PGC lines be derived (13, 14, 15). This is among the main limitation when current avian germline technologies are implemented for genetic resource preservation as well as other applications.

One reason for this bias could be related to the earlier activation of the pathways leading to the onset of meiosis in this gender compared to males; if this is correct, it would be critical that cells be isolated from embryos at the earliest possible stages (f.i. at oviposition, stage X). By doing that we run into a new difficulty: PGCs numbers in blastodermal stages are estimated in thousands (100-120). Therefore, implementing highly efficient, yet gentle procedures for PGCs isolation from blastodermal embryonic disc would be of paramount importance.

Overall studies made in Biology have focused their efforts on the protein composition, structure, expression etc. and very few have explored the role of glycoconjugates, especially glycoproteins beyond their peptide component. In general, isolation-enrichment protocols are based on monoclonal antibodies against primordial germ cell surface antigens (SSEA, EMA), which have been proven to be tedious, expensive and harmful methods (16, 9). To solve these inconvenients we propose the use of lectins, proteins that bind/recognize carbohydrates of glycopeptides. Since late XIX century, lectins were used to identify blood groups, later on it was proven that glycans were antigenic and the lectins were used to capture white cells for bone marrow transplantation in leukemia patients (17). Nowadays, again lectin microarray analysis has been used to identify and select human pluripotent cells from non-pluripotent (18). Therefore, we characterize their polysaccharide´s composition and take advantage of the distinct mucopolysaccharide profile of PGCs for their isolation, selectively binding, segregating and culturing them by means of specific lectins and their carbohydrates.

On the other hand, embryonic cells during embryo development are in a frantic movement, in fact the migration route of PGCs until they reach the gonads is well known (4). The metastazing ability of cancer cells is also related to their external mucoid envelope rich in glycoconjugates, called “glycocalix”. These mucoid envelopes are like flexible shields that confer protection to progenitor cells in their journey during development. Chemoattractant factors expressed in extracellular matrix are involved in the migration of PGCs but their capacity of migration is also related to changes in their glycoconjugate composition. Nacetylglucosamine (GlcNAc), glucosyl and mannosyl rich glycoconjugates are present in migrating PGCs but are not detected when the PGCs are established in the gonads (19). When PGCs are amplified in culture with the purpose of grafting them later onto recipient embryos to generate germ line chimerae, attention should be paid to the maintenance of a “healthy” glycocalix. Supplementation of culture media with Nacethyl D+glucosamine, one of the most abundant residues in PGCs surface oligosaccharides would facilitate the generation of the mucoid wrapping on the growing cells. To our knowledge, no such study has been done neither with PGCs nor with any other progenitor cell type.

An other distinct feature of PGCs and gonocytes, is their large size, relative to any other embryonic cell population. Yet, enrichment procedures based in cell segregation by size have not been attempted (5) so far. We developed a gentle, simple procedure for PGC isolation from stage X by size exclusion. In all our studies we consider that the albumen pH is above 8 for the first 3 days of avian embryo development, trying to emulate the alkaline environment in which avian embryos develop initially (20, 21).

Therefore, in order to improve isolation and culture conditions of PGCs, we explore in this study novel strategies for isolation of PGCs studying the external glycoprotein envelope and cellular size. We also created the conditions of a novel method for culture PGCs in which the supplementation of carbohydrate and initial alkaline culture conditions were studied. These strategies are gentle, quick and most important high quality yields of PGCs can be obtained. Potentially could be useful for many progenitor cells in Regenerative Medicine.

## RESULTS

### Characterization of PGC-LCs in stage X and HH 8-9 embryos using lectins

The pattern of sugars was very different between the two embryonic stages studied; when labelled lectin *WGA* Alexa 594 (with high affinity for N-acetylglucosamine) was used, it revealed that in stage X embryos there are cells or groups of cells clearly marked in the zone pellucida of embryo, where PGCs are localized at that stage of development. However, when *Con A* Alexa 488 (binding both D-mannosyl and D-glucosyl residues) is used, all cells are equally marked; in the centre there is no any especially marked zone (Fig. 1). On the contrary, in HH 8-9 embryos *Con A* Alexa 488, marked large cells around the cephalic area, where the PGCs are migrating at this stage of the development (Fig. 2). When *WGA* Alexa 594 is used, small and large cells are labelled over the entire embryonic body (Figs. 2, 3).

**Fig 1.**
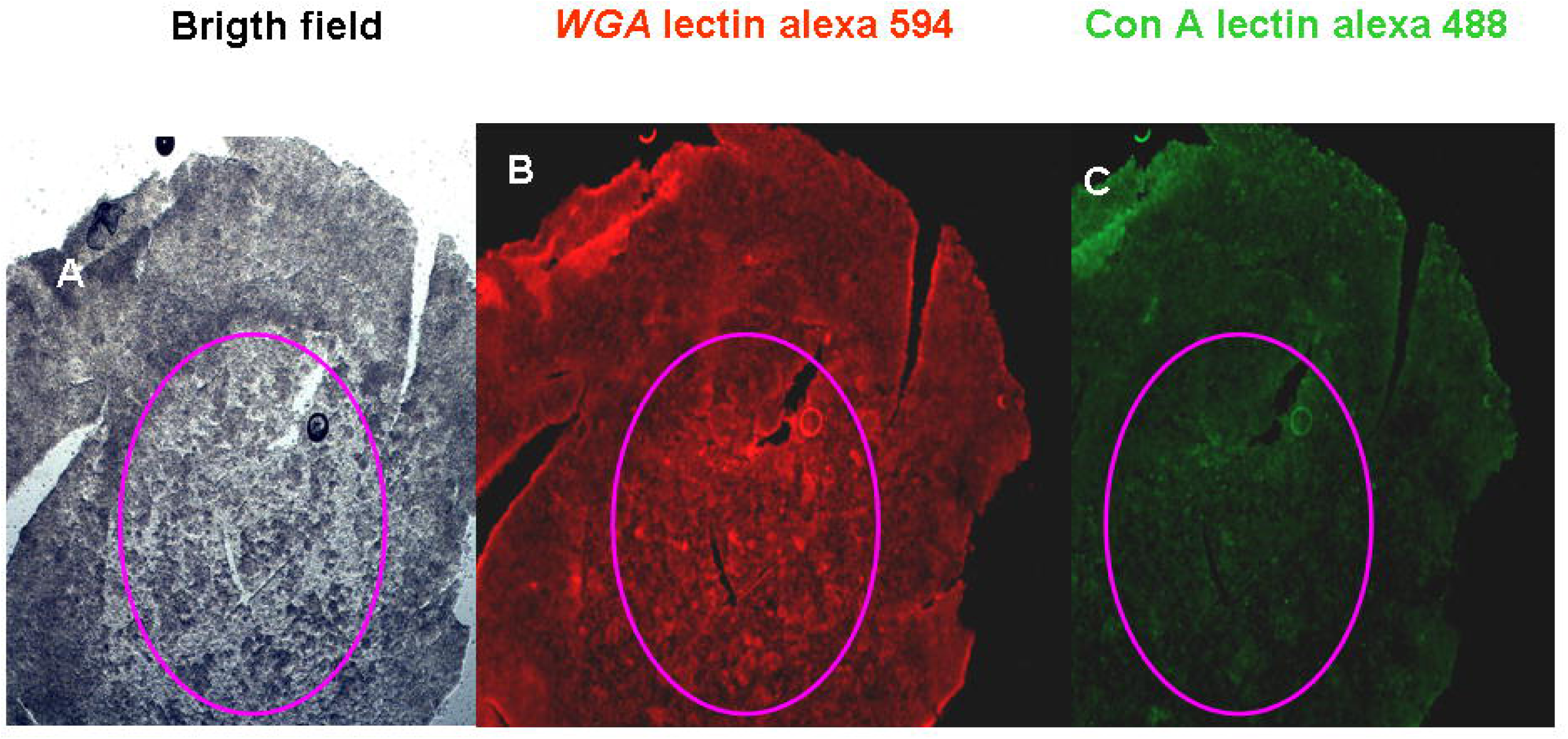
Glycohistology. Stage X embryo with *WGA* Alexa 594 and *Con A* Alexa 488. A) Bright field. B) *WGA* Alexa 594 cells marked in the zona pellucida of embryo, where the PGCs are located at this stage. C) *Con A* Alexa 488 no cells were marked in zona pellucida (pink circle) x40.

**Fig 2.**
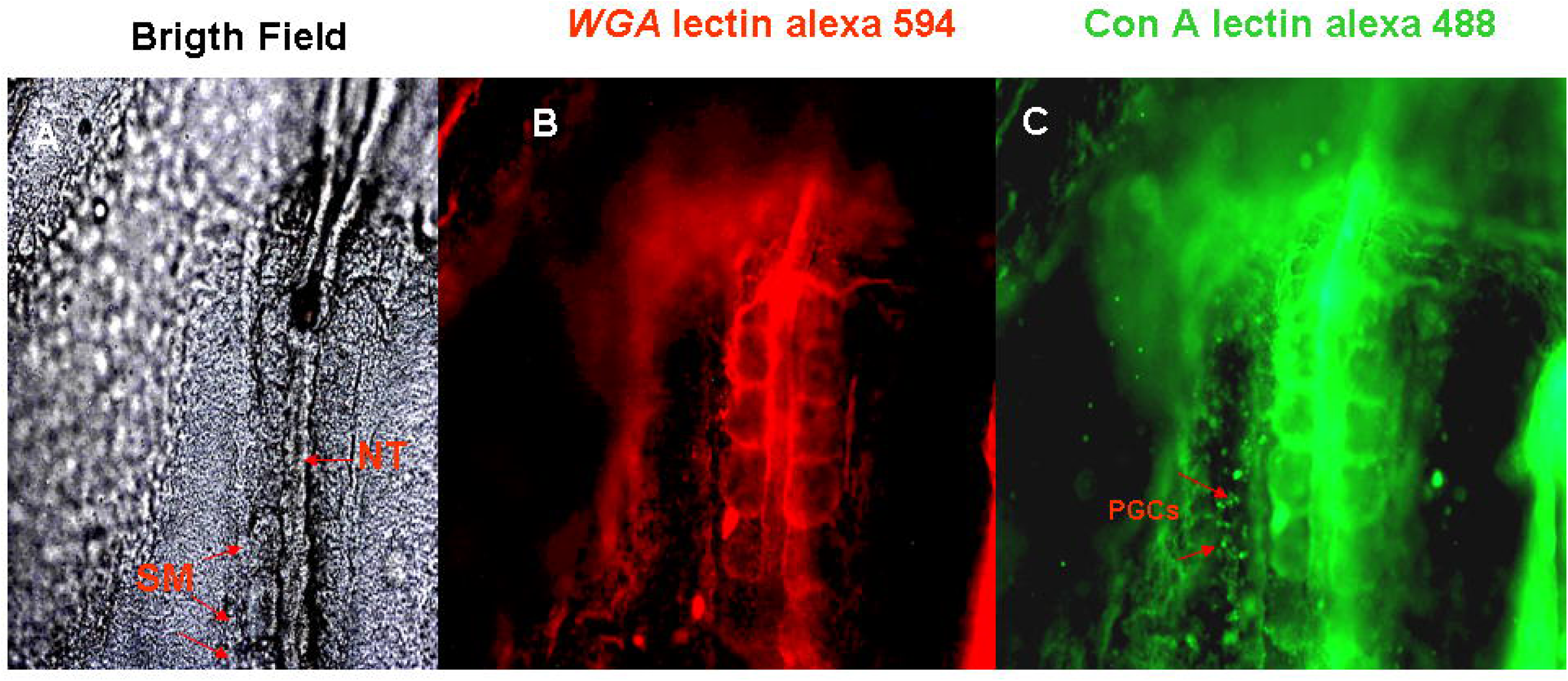
Glycohistology. HH 8-9 embryo with *WGA* Alexa 594 and *Con A* Alexa 488. A) Bright field. B) *WGA* Alexa 594 only small fluorescent cells were observed in the entire body. C) *Con A* Alexa 488 (recognizes D mannosyl residues) marked only large cells around the cephalic area, where the PGCs are migrating at this stage of development (NT notochord SM somites), x100.

Since lectins can bind to various sugars we doubted that they could reveal significant differences between the two embryonic stages. However, the pattern was not only surprisingly different but also in embryos HH 8-9 *Con A* marked large cells in the cephalic area where PGCs are expected to concentrate at this stage. To confirm that these were PGCs we conducted a double staining using lectin-glycohistology and *VASA* immunohistochemistry. It was confirmed that these large cells were PGCs (Figs. 3, 4). Therefore, lectin-glycohistology with *Con A* Alexa 488 revealed that PGCs are rich in α-D mannosyl and α-D glucosyl residues when they are migrating to the gonads (Figs. 2, 3, 4) in a specific manner.

**Fig 3.**
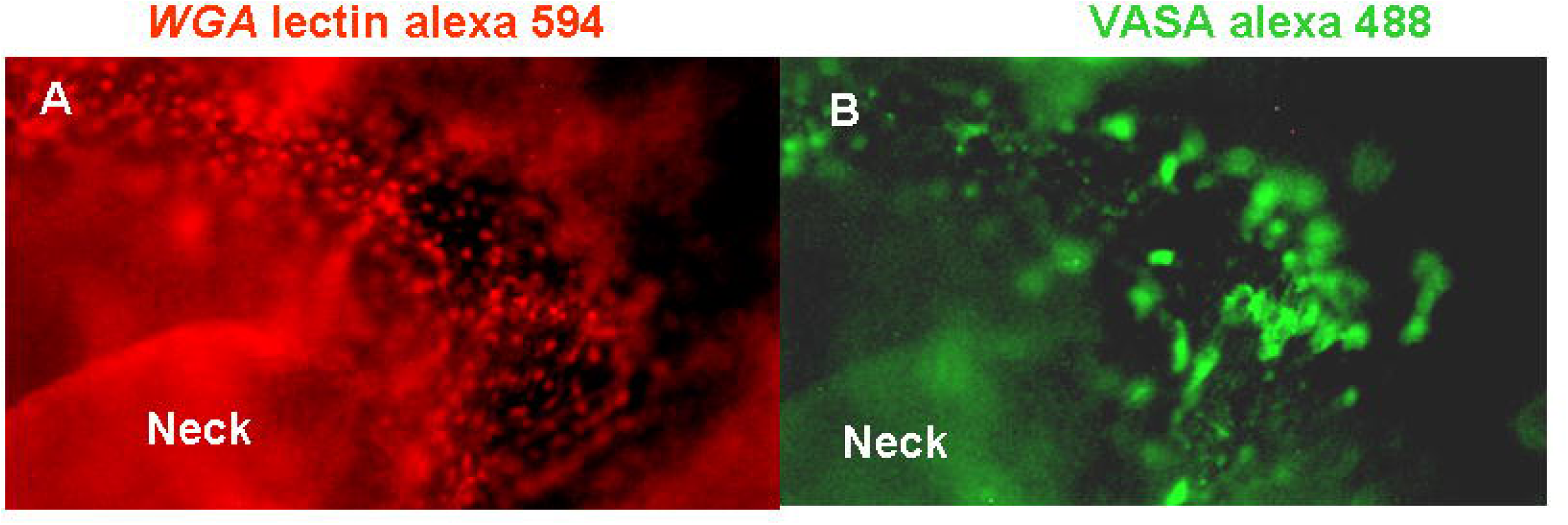
Double staining Lectin-glycohistology and immunohistochemistry of HH 8-9 embryo with *WGA* Alexa 594 and *VASA*. A) *WGA* Alexa 594 fluorescent cells were in the neck, small and large cells are marked. Large cells or PGCs are also recognized by *VASA*. B) *VASA* Alexa 488 labeled only PGCs (x100).

**Fig 4.**
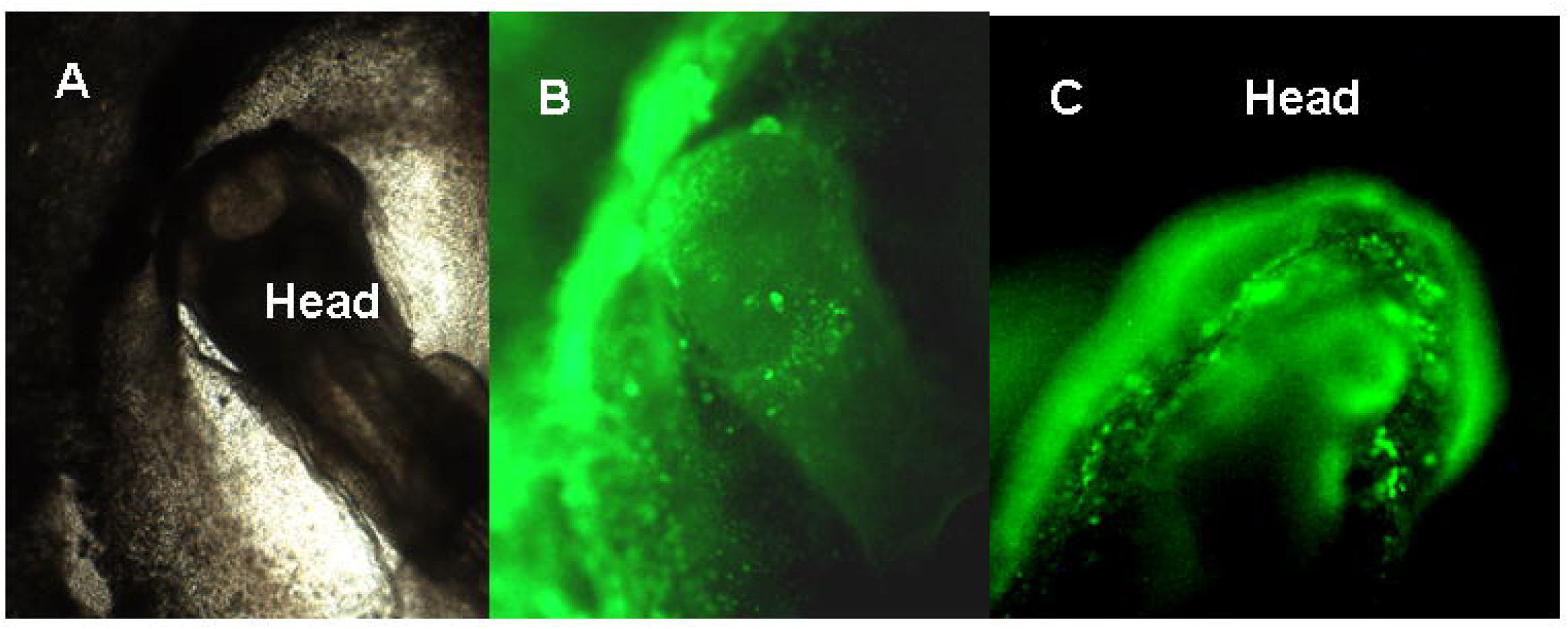
PGCs positive to VASA immunohistochemistry in cephalic area of HH 8-9 embryos. (A and B x40 and C x100).

From this study it is clear that during the embryo development the PGCs change their composition in polysaccharides. The PGCs present in stage X embryos have in their outlayer predominately Nacetylglucosamine and later on, they are rich in α-D mannosyl and α–D glucosyl residues in a specific manner.

It is important to emphasize that using labelled lectins allowed us to characterize the composition of polysaccharides in just 10 minutes, and also gave us valuable information for the next study.

### Immobilized lectins

ELISAs are based on microplates coated with antigens or antibodies (proteins) and their specific binding (antibody-antigen). Since the lectins are proteins, we considered the possibility that culture plates coated with lectins could retain cells rich in mucins as it is the case of PGCs. Based on the results obtained in the study made with labelled lectins, we coated culture plates with *WGA* and *Con A* and the binding was evaluated immediately under inverted microscope.

The uncertainty about if the affinity between lectin-glycoprotein binding was high enough was successfully unveiled immediately at the first attempt of isolation. Therefore, in the following isolations we attempted to go further and the retained cells were subjected to different culture conditions. We observed in lectin coated-plates that after one hour of incubation at room temperature, cells were loosely rooted to the plate, as if they were tethered by a transparent mucous material, and also observed that there were many cells with morphology compatible with PGCs (Fig. 5). On the other hand, it is known that the albumen that is bathing the chick embryo has a pH above 8 for the first 3 days, trying to emulate the environment in which avian embryos are developed initially (20, 21), the cells retained by lectins were cultured under an alkaline environment. Under culture conditions of alkaline pH, at 37ºC and during 48 hours both lectins were effective in retaining cells. In control plates non lectin-coated, the cells disappeared completely (Fig. 5)

**Fig 5.**
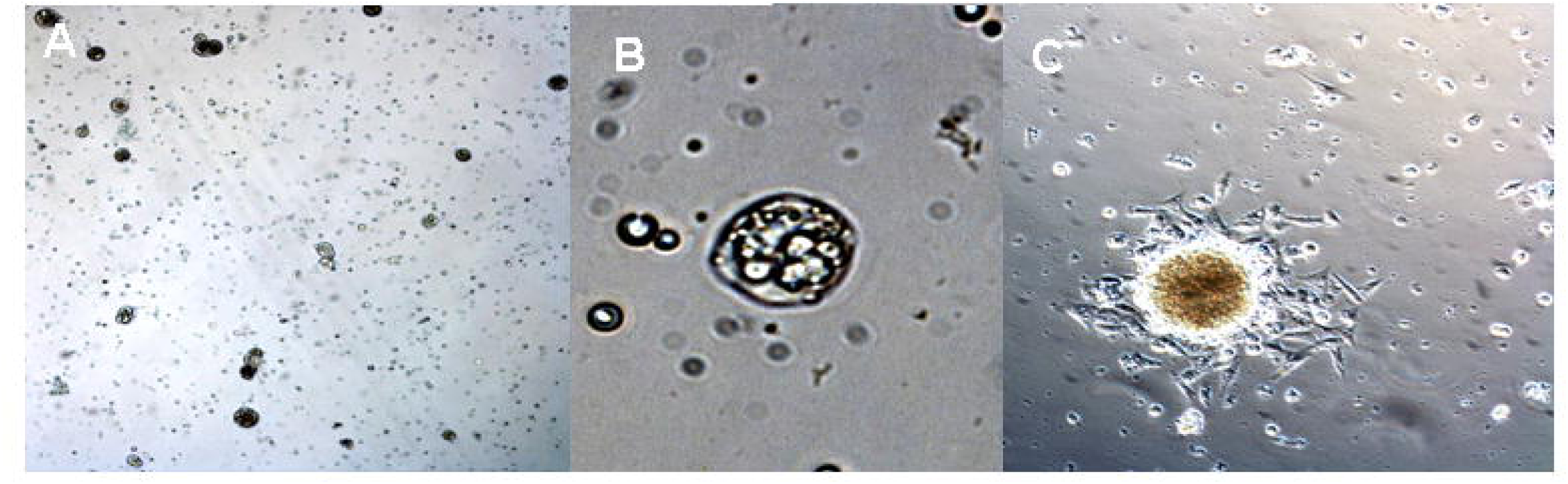
Cells isolated from Fase X and HH 8-9 embryos in coated lectins plates. Fresh cells A, B and C. A and B PGCs isolated from stage X embryos after 1 hour incubation at room temperature (x100 and x400, respectively A and B). C) Colonies of PGC isolated from HH 8-9 embryos over tightly adhered cells (x100).

The most striking results were those obtained from HH 8-9 embryos under culture conditions of alkaline pH, at 37ºC and during 48 hours. The isolated cells formed numerous clumped PGC-LCs with spherical shape, which were above a layer of cells, which in turn, were strongly bound to the plate. Some cells that were detached and floating close to the cell clusters, resembled morphologically to the PGCs previously described. Immunohistochemistry confirmed that they were PGCs and revealed that the colonies are formed by hundreds of positive cells to VASA and *SSEA-1* (Fig. 6). Cells isolated from stage X embryos attached tightly when plates were coated with *WGA*, cell retention was observed within seconds and big cells with refringent granules were plated in the first two hours under culturing conditions with alkaline pH after 48 hours at 37ºC.

**Fig 6.**
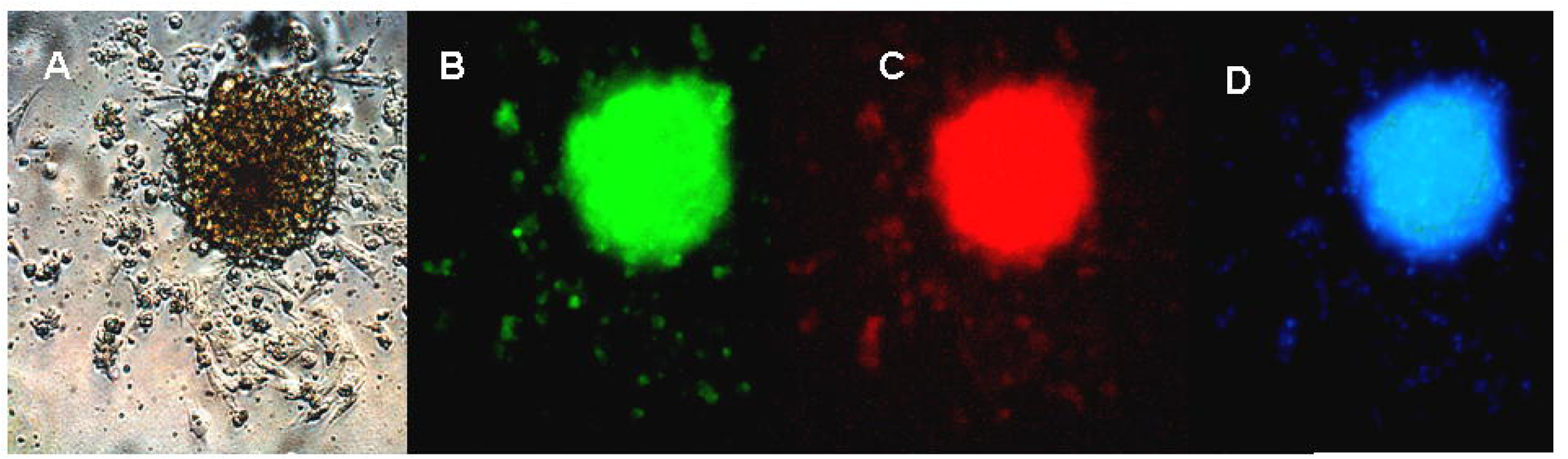
Cells isolated from HH 8-9 embryos in coated-lectin plates. Colonies of PGCs isolated from HH 8-9 embryos. Cultured under alkaline pH conditions after 48 hours. A) Bright field. B and C immunohistochemistry of *VASA* and SSEA-1, respectively. D) Hoescht (x200).

Therefore, in the plates lined with lectins the retained cells showed different morphologies. Lectins retained the PCGs and favored its cultivation when isolates from HH 8-9 embryos were used. From every single embryo seeded into coated plates thousands of PGCs were cultured in just 48 hours. Although we did not sex embryos, the same results were seen in all 15 isolates.

### Cell strainers

The bibliography is full with descriptions of PGCs and many of them agree in that they are large cells. However, during the developmental stage X although it is known they are in the zone pellucida, are very difficult to differentiate from other cells because they are scarce (100-120) and the other cells are also large (20,000-50,000). Cellular differentiation is even more complicated at the moment of isolation because there are always traces of yolk very refringent with spherical shapes. Therefore, taking advantage of the new cell strainers commercially available, that are able to discriminate particles with differences as little as of 1 μm we explored their usefulness to separate PGCs.

In stage X embryos almost half of the fresh pellet was retained in the cell strainer of 10 μm while the other half passed through the smallest size tested (6 μm estimates by pellet size). *VASA* immunohistochemistry revealed that almost all *VASA* positive cells are larger than 10 μm and the range of diameters in cells retained was 24-30 μm. On the contrary, in HH 8-9 embryos 20% of the fresh pellet was retained in the cell strainer of 10 μm the range of diameters in *VASA* positive cells retained was 14-17 μm, indicating that the PGCs are smaller at HH 8-9 versus stage X embryo (Table 1). These results indicate that the cell strainers with pore size less than 10μm can retain the majority of PGCs from stage X and HH 8-9 embryos, and most important the purity of PGCs isolated is almost complete (Fig. 7).

**Table 1.**
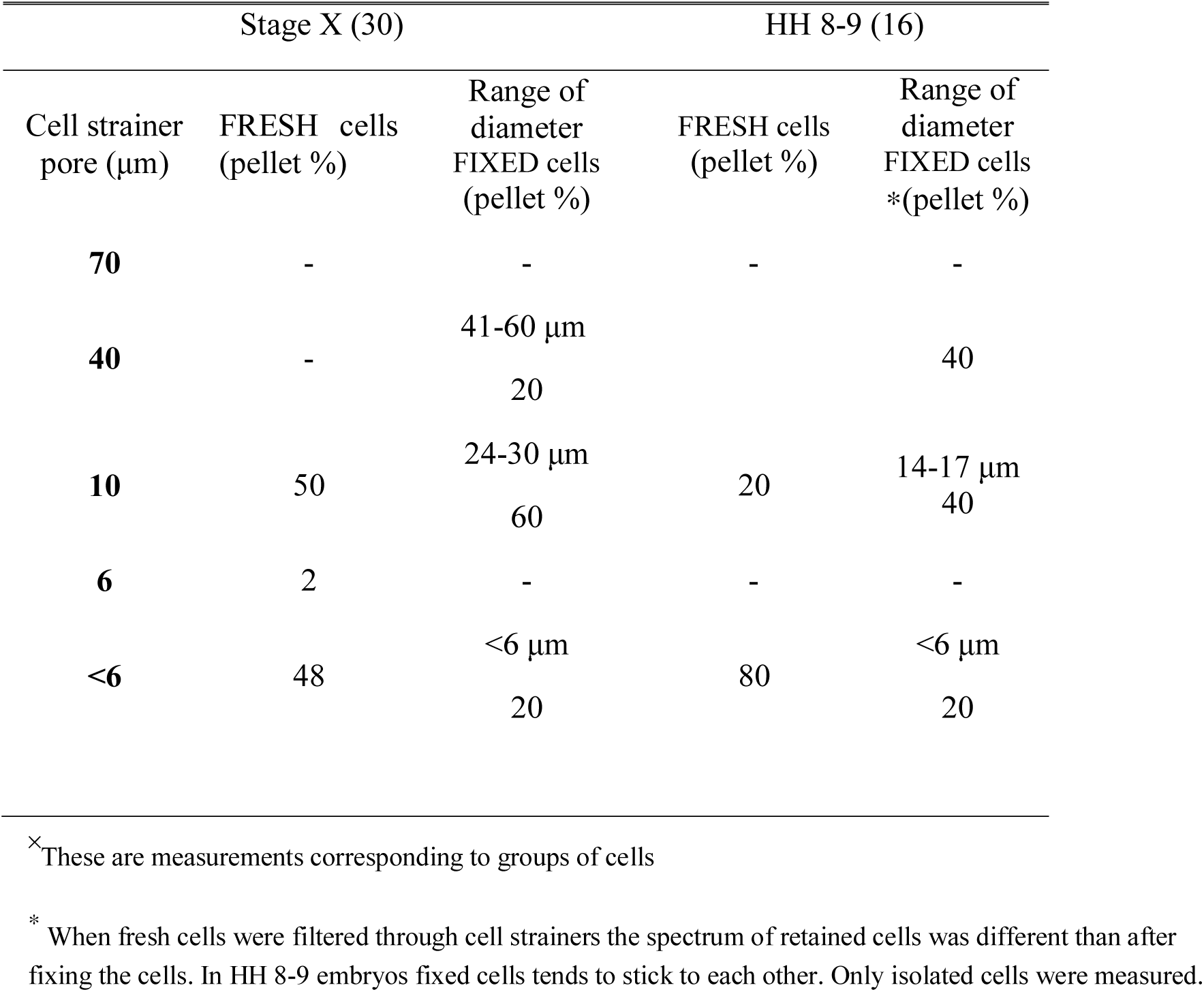
Percentage of fresh and fixed cells retained in cell strainers.

**Fig 7.**
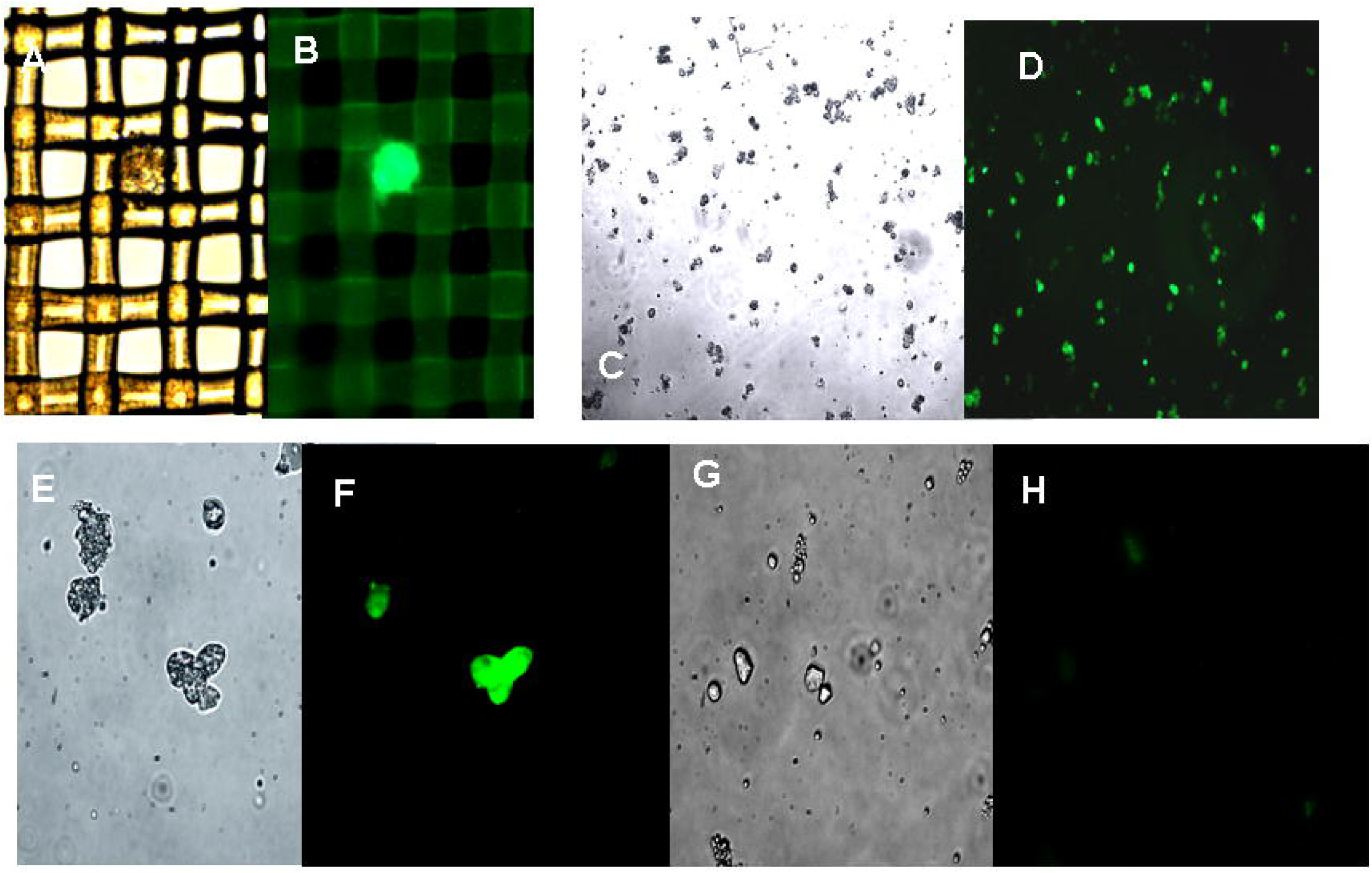
Cells isolated from Stage X embryos using cell strainer (CS). A and B cell retained in pore size of 40 μm *VASA* positive (x40); C, D, E and F cells retained in CS of 10 μm pore size are *VASA* positive (C, D and E, F are x100 and x400, respectively); G and H, cells retained in pore size of 6 μm, all are *VASA* negative (x400).

### Cultured cells

In order to obtain success in culturing, a minimum number of cells is imperative and the purity of cellular isolates is also critical. In the case of PGCs there is an additional problem: female PGCs initiate meiosis, which is incompatible with cellular division earlier than male PGCs. Most of the studies on chicken PGC isolation have been made with circulating PGCs and they have been successful with cultured male PGCs, but almost all have failed with female PGCs. Therefore, all our studies were primarily focused to obtain cultured PGCs from stage X embryos and also in stage HH 8-9 embryos because in the latter the number of cells has increased exponentially, both stages are far enough from first signals of meiosis in both gender.

PGC-LCs were seen after 4-5 days of culture of cell isolates from both stage X and HH 8-9 embryos, although the proportion was higher in stage X embryos. In stage X cultures many floating PGC-LCs were seen in all treatments used, their number being especially remarkable on the first day of culture when grown in N-acetyl-D-(+)-glucosamine supplemented germ basic medium. On day 14 of culture plenty of floating refringent cells, resembling gonocytes, were detected in this group on the bottom of the plate. Their diameter was typically larger than 30 μm, and many could be seen surrounded by a spherical capsule (26-43 μm in diameter). Some were leaving the capsule and some were mitotically active (Fig. 8).

**Fig 8.**
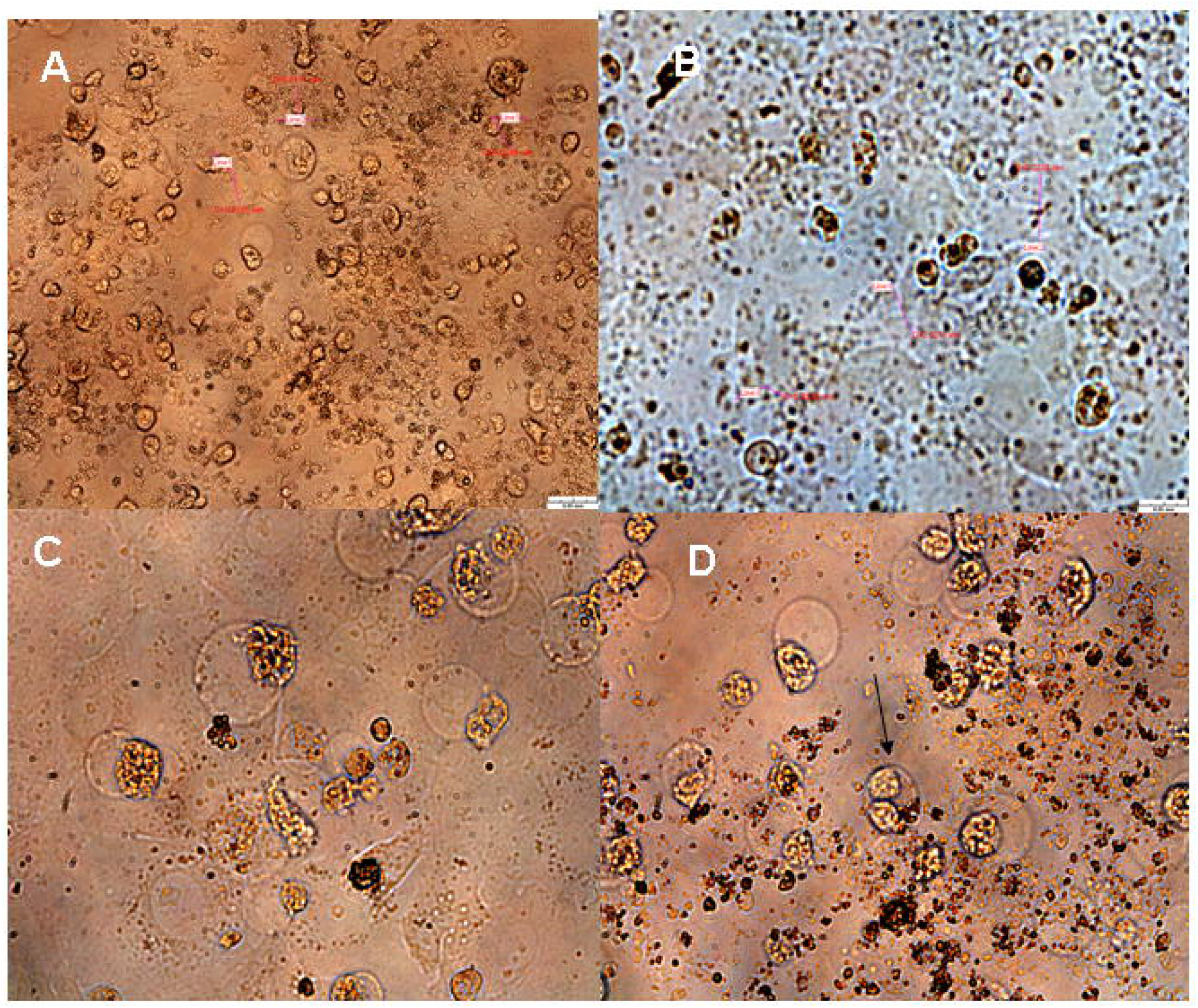
Primary culture of gonocyte-like cells in N-acethyl-D(+)-glucosamine supplemented germ cell medium. A) Cells isolated from Stage X embryo with a diameter mean of 25 μm at 17 days of culture (x200). B) Cells resembling oocyte are floating with a diameter mean of 30 μm at 14 days of culture (x200). C and D cells are very refringent, are inside an spherical capsule of 40 μm, some are leaving the capsule and other are mitotically active (black arrow) at 17 days of culture (x400).

Once the PGCs-LC started to appear, the use of hanging cell inserts allowed us an easy management of medium changes without loosing cells and a closer observation of the cellular evolution.

## DISCUSSION

In this study we demonstrate that the mucoid envelope of the PGCs in very early stages of embryo development (after oviposition, comparable to blastocyst stages in mammals) changes the glycans exposed in a specific manner as embryonic development progresses.

Our studies made for carbohydrate characterization of the PGC-LCs in stage X embryos using lectins revealed that *WGA* Alexa 594 (with high affinity for Nacetylglucosamine) labeled cells or cell clusters which could be seen all over the zona pellucida where the PGCs are located at this developmental stage. In HH 8-9 embryos we found that *Con A* alexa 488 (specific binding for αD-mannose) also provided highly efficient, specifically labeling only PGCs of whole HH 8-9 embryos, without background. In our experiments, we saw a perfect match between *Con A* alexa 488 and *VASA* labeling in HH 8-9 embryos. *VASA* is considered as the only specific antibody marker of the embryonic germline, from stage X to oocyte differentiation. Until now good antibodies against chicken *VASA* were limited but the one used in this study turned out to be very specific and commercially available. Therefore, in HH 8-9 embryos *Con A* alexa 488 is specific and can be used alone as marker or as a capturer. However, we must emphasize that lectin histochemistry advantages are numerous over immunohistochemistry; particularly it is more stable, cheaper and faster (one single step assay, reaching a clear signal in a few minutes, with negligible background). Above all of these advantages, lectins are better characterized with respect to binding specificity than monoclonal antibodies.

Our studies indicate that a progressive change in the carbohydrate profile is happening after oviposition, taken into consideration the spatiotemporal dynamics of PGCs and the results using lectin and immune labeling procedures. We did not expect lectins to have such specificity for PGCs in such a narrow window of time. Cells in blastodisc stage X are less differentiated than in HH 8-9 embryos and we know that the less differentiated a cell is; the less distinct transcriptional factors are expressed. In fact, in stage X embryos it is very difficult to differentiate PGCs from other cells using standard histological procedures (PAS, haematoxilin eosin, etc). However, *WGA* and *Con A* marked PGCs with high specificity in stage X and with an exquisite specificity and affinity in HH 8-9 embryos, respectively.

Moreover, we also show that lectins selectively retain PGCs on culture dishes lined with these proteins. In our first attempt, lining for 1 hour at room temperature was enough to immobilize stage X embryo PGC-LCs in a matter of seconds after seeding. It has been reported that *Kds* for lectins-monosaccharides binding in ELLA (enzyme lectin labeled assays) ranges between 2⨯10^−6^⨯10^−7^ M (22); we demonstrated that these *kds* are strong enough to retain PGCs. Further studies on lectin-coated plates would allow the optimization of the plate coating conditions (temperatures, coating time, lectin concentration, reaction time, etc). The coated-lectins plates could be very well used in kinetic studies made with valuable cell lines (f. i. fetal human cell lines) to figure out the lectin with the highest *Kd* and even to characterize the α or β sugar structure of the glycoconjugate(s) with binding competitive studies using different haptens.

Because of the success obtained in the number of PGC-LCs retained, in our next experiments we tried to culture the bound PGCs-LC with germ basic culture medium emulating the alkaline conditions of the blastodisc within the egg the first three days (22). Under these conditions, in just one step we were able to isolate and culture thousands of PGCs forming numerous germspheres in 48 hours from one single HH 8-9 embryo. Later on, PGCs were further amplified and subcultured under standard coculture conditions (13, 23, 24). As expected, given the PGC count at different stages and assuming similar lectins´s *Kds* in all stages, the number of retained PGCs was higher in HH 8-9 than in stage X embryos. We must emphasize that not only the number of PGCs recovered per embryo was remarkable, but also we had same efficiency in all isolates. The large number of harvested PGCs would allow further manipulations like transgenesis or germline chimera construction. For instance, female germline chimeras could be produced with an expected 50% success, after PCR-sexing of these cultures, but not the receptor embryos.

Despite the differences observed between lectins in our PGC characterization studies, both, *Con A* and *WGA*, were effective in retaining PGCs on the plates. This might be due to the cross-reactivity of lectin with several sugars (*Con A* binds αDmannose but also αDglucose). Also, we did not exhaustively count the number of cells bound for each ligand; therefore, differences might exist that were not quantified.

We also succeeded in segregating, with a very high purity, PGCs from stage X and HH 8-9 gross cell suspensions through cell strainers (size exclusion > 10μm), characterizing them morphometrically (>15 μm in diameter for HH 8-9 embryos and even larger for stage X >20 μm, as expected) and immunohistochemically (*VASA*). Up to date, the cell strainers commercially available had as minimum a pore size of 40 μm, however, we used new commercially available cell strainers of 30, 20, 10 and 6 μm pore size, which allowed us a very fine cell sieving.

The isolation efficiency of stem cells is an important challenge in Regenerative Medicine. In general, target progenitor cells are scarce and disperse, making their isolation difficult from almost any tissue. For their successful culture, a minimum number of cells is imperative and the purity of cellular isolates is also critical. As in the present study, labeled lectins could be used for embryonic stem cell differentiation studies or for isolation purposes from stem cell niches in differentiated tissues. Surface carbohydrates of the target cell population should be firstly characterized “in situ” with labeled lectins. The appropriate raw lectin and the hapten can then be used to trap and enrich these cells on culture dishes, respectively. Likewise, cell strainers could be used as an easy and gentle procedure for the segregation of specific cell lineages from tissue homogenates of different origins.

In terms of PGC-LCs long-term culture, we were able to maintain and grow PGCs and gonocyte like cells (mitotically active) from stage X embryos for 20 days in culture. We have not found any study or data with these strategies showing such efficiency from stage X embryos. This advantage is especially important if we want to culture female PGCs, because the farther in time the signals of the meiotic phase are, the better. It has been proven that cultured PGCs and gonocytes are able to colonize the gonads and that the efficiency of germline transmission is higher when PGCs are sourced from less developed embryos (24). The earliest source used by the majority of studies in chicken is circulating PGCs because, although they are not completely naïve, they are present in blood at relatively high concentrations and handling them in a liquid matrix is easier. These cells have been successfully amplified in vitro, but, after long periods in culture, two significant problems arise: a clear bias for male PGCs, and a substantial loss of migration capacity (11). Although we did not test the gonad colonization and germ line transmission ability of our cells, this capability has been reported to be lost after longer periods in culture (11, 12) than in our experiments (77-111 days, vs. 17-20 days in ours).

We evinced that N-acethyl D+glucosamine supplementation to the culture media could greatly enhance the amplification of isolated PGC and that gonocyte like cells were very well maintained in culture until day 20. Supporting our results are the reported results obtained in the mammalian nervous system, also rich in glycans, in which the single addition of GlcNAc initiates the biosynthesis of complex-type N-glycans which cover neural progenitor cells NPCs (25). The glycocalix of adult NPCs allowed to Hamanoue et al. to develop a lectin panning method based in lectin-coated plates similar to ours (26, 27, 28). Very few studies have been made in which lectins were used to isolate from adult tissues either precursor lymphocyte T (17) or NPCs (26). However, the differentiating feature in our study is the enrichment procedure, once the cells were trapped by lectin we add the carbohydrate, while in these other studies the authors tried to favour the cell culture conditions adding more lectin. Moreover, supporting our results in a very recent study Hamanoue has suggested that the glycans (biantennary and β1, 6-branched N-glycans) and their enzymes
(N-acetylglucosaminyltransferase GnTV) are implicated in NPCs migration and proliferation, rather than cell–cell attachment (29). We suggest that carbohydrates of glycans covering progenitor cells are also implicated into cell multiplication and most probably in cell migration and protection.

Interestingly, N-acethyl D+glucosamine is a structural component of microbial walls commonly used in microbiological media, it is also present in chicken PGCs and mouse NPCs. Maybe, this very well conserved carbohydrate residue from bacteria to eukaryotes cells can also share same immunological protective functions. We propose that cells cultured in vitro supplemented with the adequate carbohydrate could have an important role in cellular immunity protection, preventing the rejection of stem cells transplantation. Including as a supplement not only in chicken PGC growth media but also in other progenitor cell media like NPCs medium, could be the source for important glycoproteins and glycolipids. The very idea that a proper chemically identification of the carbohydrate and its addition for “in vitro” production of unrejectable mammalian stem and primordial germ cells it is a fascinating challenge.

Paraphrasing Cummings, “The lectins specificity has catapult the field of glycobiology into the modern era”. In our opinion, in view of our results together with the fact that lectins can differentiate between α or β anomers of carbohydrates in glycans (3D sugar chemical structure) could catapult the field of glycobiology beyond the regenerative medicine” (22).

In summary, we have first shown that N-acetyl D+glucosamine is an abundant constituent of the outer cell membrane in the earliest developmental embryonic stages; second that binding by specific lectins is strong enough to isolate and retain PGCs and, third it is an appropriate supplement for long-term PGCs culture. These three propositions configure an argumental strength supporting the crucial role of carbohydrates of glycoconjugates in PGCs culture.

Nowadays, the real possibility to insert and control the gene expression in germ cells is crucial, because it guarantees stable transmission of the transgene to the germline. The knowledge of the chicken genome together with the development in genetic engineering to insert genes into certain cells or whole animals (30) predict extraordinary advances in next years ahead. Enthusiastically, we expect that the procedures exposed in this study will help to make important advances in regenerative medicine.

## CONCLUSIONS

Herein we reported highly efficient, yet gentle procedures for PGCs isolation from blastodermal embryonic. The novel strategies improved the efficiency of PGCs isolation taking advantage of two distinct features of the early PGCs: their large size and their unique mucopeptide envelope. A rapid enrichment is obtained by one step screening using the appropriate cell strainers. On the other hand, a panning procedure with lectin coated culture dishes allows a selective retention. Followed by N-acethyl D+glucosamine supplementation to the culture media which enhanced the multiplication of PGCs.

The isolation and culture strategies developed for embryonic PGCs, could also be applied to a range of other progenitor cells present in very low numbers but with great interest for stem cell manipulations and for regenerative medicine applications.

## METHODS

### Egg source

Fertilized eggs were purchased from local suppliers in San Diego (CA, USA) that regularly supply the Salk Institute for research purposes.

### Stage X and HH 8-9 embryos isolation

Embryos in stage X were dissected after dipping the yolks into cold saline solution. Fertilized eggs were incubated until chicken embryos reached stages HH 8-9; the cranial part of the embryos was dissected by using discs of filter paper because at these stages most of the PGCs are in the cephalic area. After dissection, the embryos were cleaned free from yolk residues and vitelline membranes under stereoscope. All isolations and some initial culturing conditions of PGCs were made under alkaline environment (21) and later under standard culture conditions (13, 24).

### Cell isolations

The cells were dispersed by repeated pipetting, no enzymes were used. Two isolation procedures were tested in disaggregated cells from stage X and HH 8-9 embryos, some were seeded onto lectins-coated plates with *wheat germ agglutinin* (*WGA*) or *Concanavalin A (Con A)* and others filtered through six cell-strainers of decreasing pore size. Histochemical studies were made on PFA-fixed embryos “in toto”, and on isolated cells either in fresh or after fixation with 5% PFA.

### Characterization of Stage X and HH 8-9 embryos and cell isolates using lectins

*Lectin-histochemistry.* Two labeled lectins, *WGA-Alexa 594* and *Con A-Alexa 488* (Lifetechnologies), were used at manufacturer’s recommended working conditions. Fixed stage X and HH 8-9 embryos were blocked with 3% BSA and 0.1% triton X-100 in PBS overnight at 4ºC. Later on, embryos “in toto” were incubated with labeled lectins at 37ºC for 10 minutes. Also labeled lectins were used for the characterization of isolated cells with immobilized lectins.

2.4.2. *Immobilized lectins* were used to capture PGCs from stage X and HH 8-9 embryos (10 and 6, respectively) on 35 mm culture dishes (Falcon) coated with *WGA* (high affinity for *N*-acetylglucosamine) or *Con A* (binding both D-mannosyl and D-glucosyl residues) at a concentration of 250 mg/ml for 1.5 hours at room temperature. Excess of lectins was discarded and the plates washed with DPBS. Dispersed cells from stage X and HH 8-9 embryos were incubated for two days without CO_2_ at 37º C; under these conditions the medium reaches alkaline pH values emulating albumen pH first three days of embryonic disc development (20, 21). After that, cells were detached from the plates using specific dilutors for both lectins with 500 mM *N*-acetylglucosamine (Vector Labs ES-5100) and 200 mM α-methylmannoside/200 mM α-methylglucoside (Vector Labs ES-1100) for *WGA* and *Con A* coated plates, respectively. On the third day the eluted cells were seeded into 48-well plates and co-cultured with inactivated BRL under standard conditions with 5% CO_2_, at 37ºC (24).

### Cell strainers

A total of 30 stage X embryos (three batches, 10 embryos/batch) and 15 embryos at stages HH 8-9 (three batches, 5 embryos/batch) were filtered through cell strainers. The cell mass recovered from each strainer was evaluated under stereoscope and inverted microscope. In order to assert that sieving was working, measurements of filtered cell diameters were taken. Cell strainers were stacked from up the larger pore size (70 μm) on top, down to the smallest (6 μm). Pore sizes used were 70 μm, 40 μm (Falcon), 30 μm, 20 μm, 10 μm and 6 μm (Pluriselect). The mass of filtered cells was recovered from each cell strainer and centrifuged at 200xg for 15 minutes. Each pellet recovered was assigned a percentage such as the sum of all estimated pellet were 100%.

Immunohistochemistry with *VASA* antibody was made in order to identify where the PGCs were retained. Since it was proven that the majority of the *VASA*-positive cells was found to be retained in cell strainers of 10 μm, we decide to use 24-well plates co-cultured with feeder BRL cells using hanging cells inserts of 5 μm (as minimum twice smaller than cells diameter; Millicell Hanging Inserts Millipore). This allows us a closer follow up of PGCs evolution.

### Co-Culture conditions

*Culture media*. Germ cell culture medium used was the same as described in others works (van de Lavoir et al. 2006a; Song et al. 2014 and Miyahara et al. 2014). The medium consisted of knockout DMEM (Invitrogen), 40% buffalo rat liver (BRL)-conditioned KO DMEM, 7.5% FBS (Standard Hyclone), 2.5% Chicken Serum (Sigma), 2 mM GlutaMax (Invitrogen), 1 mM sodium pyruvate (Invitrogen), 1xNEAA (Invitrogen), 0.1 mM β-mercaptoethanol (Invitrogen), 4 ng/ml recombinant human FGF basic (rhFGFb) (R&D Systems), 6 ng/ml recombinant murine SCF (rmSCF) (R&D Systems), 1% Pen-Strep (Invitrogen). Sera used were previously frozen at −20ºC, but not inactivated by heat. Complement inactivation is achieved if sera are frozen at higher temperatures of −70ºC (Mariacruz López Díaz, patent P2001431678). Feeder BRL cells (ATTC 1442) were expanded and seeded at a concentration of 10^5^cells/ml in 48, 24 and 12-wells plates. When BRL cells reached 70% of confluence were inactivated with mytomicin 10mg/ml, one day before adding embryo cells.

Cells from each embryo were separately seeded in 48-wells plates coated with matrigel and with a feeder layer of mytomicin treated BRL cells. In all plates controls wells were seeded with cells from HH 8-9 embryos following Song et al. procedure (2014). Cells from stage X embryos were seeded after either filtering through cell strainers (> 10 μm), or isolation with lectins and their dilutors, or with no previous fractionation.

Sixteen wells were seeded with cells from stage X embryos and cultured in germ cell medium supplemented N-acethyl-D(+)-glucosamine (Sigma; 0,3 mg/ml).

After a week when numerous PGC-LCs were seen in suspension, approximately every week cell cultures were passed to larger well plates; first to 24-well plates using hanging cells inserts of 5 μm, and then to 12 well plates without inserts. Hanging cell inserts of that size guaranteed a fluid media exchange and prevented loosing cells.

### Characterization of PGC-LCs

*Immunohistochemistry of PGC-LCs in cellular isolates and embryos.* PGC-LCs were identified in all embryos and cell isolates by immunohistochemistry and glycoprotein histochemistry.

Anti-*VASA* ( H-80 Cat#: sc-67185 Santa Cruz Biotechnology rabbit IgG polyclonal) was used for immunohistochemistry since *VASA* gene is expressed only in germ cells and gonocytes at all stages of development, and also anti-*SSEA-1* (Hybridoma bank monoclonal antibody). Fixation was performed in 5% PFA in PBS for 15 minutes at RT (isolated cells). To minimize nonspecific binding, the fixed cells and embryos were treated for 3 hours with 3% BSA and 0.1% triton X-100 in PBS before immunostaining. Embryos and cells were incubated for 24 hours with the primary antibodies at 4ºC and subsequently reacted for 24 hours each with alexa-568 donkey anti-mouse (for monoclonal first antibody, Invitrogen) and alexa-488 donkey anti-rabbit (for polyclonal first antibody, Invitrogen). The optimal concentration of each antibody was selected based on the recommended manufacturer’s conditions for primary antibodies (1/30 and 1/50 for anti-VASA and anti-SSEA1, respectively) and 1/500 for both secondary antibodies.

## DECLARATIONS

### List of Abbreviations

PGCs: Primordial germ cells
ESCs: Embryonic stem cells
iPSC: induced pluripotent stem cells
IMACS: Magnetic-Activated Cell Sorting
FACS: Fluorescent-Activated Cell Sorting
SSEA1: Stage-specific embryonic antigen-1
EMA: Primordial germ cell marker (mouse)
WGA: Wheat germ agglutinin
Con A: Concanavalin A
PGC-LC: Primordial germ cell like cell
PFA: Paraformaldehyde
PBS: Phosphate buffer saline
DPBS: Dulbecco phosphate buffer saline
BRL: Buffalo Rat Liver
FBS: Fetal bovine serum
DMEM-KO: Dulbecco modified eagle medium knockout
rhFGFb: recombinant human FGF basic
rmSCF: recombinant murine SCF
NPC: Neural progenitor cell
GlcNAc: N-acethyl D+glucosamine
GnTV: N-acetylglucosaminyltransferase

**Ethics approval and consent to participate** ‘Not applicable’

**Consent for publication** ‘Not applicable’

**Authors’ contributions** ‘Not applicable’

## Availability of data and material

The datasets during and/or analysed during the current study available from the corresponding author on reasonable request.

## Competing interests

The authors declare no competing or financial interests.

## Founding source

This study was undertaken thanks to all resources available at Gene Expression Laboratory, Salk Institute for Biological Studies, 10010 North Torrey Pines Road, La Jolla, California 92037, USA. Sponsored by Dr Juan Carlos Izpisua Belmonte. Also this study was partly supported by the grant AGL 2009E06345–MICINN (Spain).

## Acknowledgements

This study could not be undertaken without the resources and hospitality of Dr Juan Carlos Izpisua Belmonte´s Lab at Salk Institute (La Jolla San Diego, CA).

The author acknowledges the valuable support of Dra Capitolina Díaz Martínez President of AMIT (Asociación de Mujeres Investigadoras y Tecnólogas). Also thanks to the Direction of INIA (Instituto Nacional de Investigaciones y Tecnologías Agraria y Alimentaria). This study was partly supported by the grant AGL 2009E06345–MICINN (Spain).

The monoclonal antibodies anti-SSEA-1 developed by Solter and Knowles, and Spradling, and Williams, respectively, were obtained from the Developmental Studies Hybridoma Bank developed under the auspices of the NICHD and maintained by The University of Iowa, Department of Biology, Iowa City, IA 52242.

## REFERENCES

[1] EsfandiariE., MashinchianO., AshtianiM. K., GhanianM. H., HayashiK., SaeiA.A., MahmoudiM., BaharvandH. (2015). Possibilities in Germ Cell Research: An Engineering Insight Trends Biotechnol. Dec;33(12):735–46. doi:10.1016/j.tibtech.2015.09.004. Epub 2015 Oct 20.

[2] SugawaF., Araúzo-BravoM. J., YoonJ., KimK. P., AramakiS., WuG., StehlingM., PsathakiO. E., HübnerK., SchölerH. R. (2015). Human primordial germ cell commitment in vitro associates with a unique PRDM14 expression profile. EMBO J. 2015 Apr 15;34(8):1009–24. doi:10.15252/embj.201488049. Epub 2015 Mar 6.

[3] Dubois, R. (1969). Données nouvelles sur la localisation des cellules germinales primordiales dans le germe non incubé de poulet. C. R. Acad. Sci. 269, 205–208.

[4] Kuwana, T. and Fujimoto, T. (1984). Locomotion and scanning electron microscopic observations of primordial germ cells from the embryonic chick blood in vitro. Anat. Rec. 209, 337–343.

[5] Muniesa, P. and Dominguez, L. (1990). A morphological study of primordial germ cells at pregastrular stages in the chick embryo. Cell Differ Dev. Aug; 31(2):105–17.

[6] Swift, H. M. (1914). Origin and early history of the primordial germ-cells in the chick. Am J Anat. U S A. 15:486.

[7] Yasuda, Y., Tajima, A., Fujimoto, T. and Kuwana, T. (1992). A method to obtain avian germ-line chimaeras using isolated primordial germ cells. J. Reprod. Fert. 96, 521–528.

[8] Park, T. S., Jeong, D. K., Kim, J. N., Song, G. H., Hong, Y. H., Lim, J. M., Han, J. Y.(2003). Improved germline transmission in chicken chimeras produced by transplantation of gonadal primordial germ cells into recipient embryos. Biol Reprod. May; 68(5):1657–62.

[9] Mozdziak, PE, Angerman-Stewart, J., Rushton, B., Pardue, S. L., Petitte, J. N. (2005). Improved Isolation of chicken primordial germ cells using fluorescence-activated cell sorting. Poult Sci Reprod. 84(4):594–600.

[10] MacdonaldJ., GloverJ. D., TaylorL., SangH. M., McGrewM. J. (2010). Characterisation and germline transmission of cultured avian primordial germ cells. PLoS One. 2010 Nov 29;5(11):e15518. doi:10.1371/journal.pone.0015518.

[11] Miyahara, D., Mori, T., Makino, R., Nakamura, Y, Oishi, I, Ono, T., Nirasawa, K., Tagami, T and Kagami, H. (2014). Culture conditions for maintain propagation, long-term survival and germline transmission of chicken primordial germ cell-like cells. J. Poult. Sci. 51: 87–95.

[12] MiyaharaD., OishiI., MakinoR., KurumisawaN., NakayaR., OnoT., KagamiH., TagamiT. (2016). Chicken stem cell factor enhances primordial germ cell proliferation cooperatively with fibroblast growth factor 2. J Reprod Dev. Apr 22;62(2):143–9. doi:10.1262/jrd.2015-128. Epub 2015 Dec 2.

[13] van de Lavoir, M. C., Diamond, J. H., Leighton, P. A., Mather-Love, C., Heyer, B. S., Bradshaw, R., Kerchner, A., Hooi, L. T., Gessaro, T. M., Swanberg, S. E., Delany, M. E. and Etches, R. J. (2006a). Germline transmission of genetically modified primordial germ cells. Nature 441, 766–769.

[14] NaitoM., HarumiT., KuwanaT. (2015). Long-term culture of chicken primordial germ cells isolated from embryonic blood and production of germline chimaeric chickens. Anim Reprod Sci.Feb; 153:50–61. doi:10.1016/j.anireprosci.2014.12.003. Epub 2014 Dec 24.

[15] Whyte, J. Glover, J.; Woodcock, M.; Brzeszczynska, J.; Taylor, L.; Sherman, A.; Kaiser, P.; McGrew, M. (2015). FGF, Insulin, and SMAD Signaling Cooperate for Avian Primordial Germ Cell Self-Renewal. In: Stem Cell Reports, 01.11.

[16] EtchesR.Methods for selecting germ-line competent cells in chicken embryos, and the use of the cells in the production of chimeras. Sep 1998. International Patent Classification. C12N 5/06, A01K 67/027.

[17] Reisner, Y. and Sharon, N. (1984). Fractionation of subpopulations of mouse and human lymphocytes by peanut agglutinin or soybean agglutinin. Methods Enzymol. 108:168–79.

[18] Sancho-Martinez, I., Nivet, E., Izpisua Belmonte, J. C. (2011). Purging and isolating pluripotent cells, "sweet" dreams become true?. Cell Res; 21(11):1526–7.

[19] Cuminge, D. and Dubois, R. (1974). Role of glycoproteins in chemotactic migration of chick germinal cells, results with concanavalin A in organ culture. C R Acad Sci Hebd Seances Acad Sci D. Sep 16; 279(12):99.

[20] Stern, C.D. (1991). The sub-embryonic fluid of the egg of the domestic fowl and its relationship to the early development of the embryo. In: Avian incubation. (ed. S.G. Tullett) London: Butterworths. pp. 81–90.

[21] Lopez-Diaz, MC, Bujan-Varela, J., Cadorniga-Valiño, C. (2016). Viable pluripotent chick blastodermal cells can be maintained long term in an alkaline defined medium. In Vitro Cellular & Developmental Biology - Animal (9989). DOI:10.1007/s11626-015-9989-5.

[22] Cummings, R. D. and Etzler, M. E. In: Varki A, Cummings RD, Esko JD, Freeze HH, Stanley P, Bertozzi CR, Hart GW, Etzler ME, editors. Essentials of Glycobiology. 2nd edition. Cold Spring Harbor (NY): Cold Spring Harbor Laboratory Press; 2009. Chapter 42.

[23] van de Lavoir, M. C. and Mather-Love, C. (2006) Avian Embryonic Stem Cells. Methods in Enzymology. 418, 38–64.

[24] Song, Y., Duraisamy, S., Ali, J., Kizhakkayil, J., Jacob, V., Mohammed, M. A., Mohammed A. Eltigani, M. A., Amisetty, S., Shukla, M. K., Etches, R. J.and van de Lavoir, M-C., (2014). Characteristics of Long-Term Cultures of Avian Primordial Germ Cells and Gonocytes. Biol Reprod. 90(1):15, 1–8.

[25] Kleene, R. and Schachner, M. (2004). Glycans and neural cell interactions. Nat Rev Neurosci. Mar; 5(3):195–208.

[26] Hamanoue, M., Sato, K., Takamatsu, K. (2008). Lectin panning method: the prospective isolation of mouse neural progenitor cells by the attachment of cell surface N-glycans to Phaseolus vulgaris erythroagglutinating lectin-coated dishes. Neuroscience. Dec 10; 157(4):762–71.

[27] Hamanoue, M., Matsuzaki, Y., Sato, K., Okano, H. J., Shibata, S., Sato, I., Suzuki, S., Ogawara, M., Takamatsu, K., Okano, H. (2009). Cell surface N-glycans mediated isolation of mouse neural stem cells. J Neurochem. Sep; 110(5):1575–84.

[28] Hamanoue, M. and Okano, H. (2011). Cell surface N-glycans-mediated isolation of mouse neural stem cells. J Cell Physiol. Jun; 226(6):1433–8.

[29] Hamanoue, M., Ikeda, Y., Ogata, T., Takamatsu, K. (2015). Predominant expression of N-acetylglucosaminyltransferase V (GnT-V) in neural stem/progenitor cells. Stem Cell Res. Jan; 14(1):68–78.

[30] Lillico, S.G., Sherman, A., McGrew, M. J., RobertsonC. D., Smith, J., Haslam, C., Barnard, P., Radcliffe, P. A., Mitrophanous, K. A., Elliot, E. A., Sang, H. M. (2007). Oviduct-specific expression of two therapeutic proteins in transgenic hens. Proc Natl Acad Sci U S A. Feb 6; 104 (6):1771–6.

